# Do camera light traps for moths provide similar data as conventional funnel light traps?

**DOI:** 10.1101/2025.02.06.636905

**Authors:** Vivian Holzhauer, Dennis Böttger, Paul Bodesheim, Gunnar Brehm

**Affiliations:** Institut für Zoologie und Evolutionsforschung mit Phyletischem Museum, Friedrich-Schiller Universität Jena, Jena, Germany; Biologische Station Lippe e.V., Domäne 2, Schieder-Schwalenberg, Germany; Computer Vision Group, Friedrich Schiller University Jena, Jena, Germany

**Keywords:** arthropods, conservation, insects, Lepidoptera, monitoring, camera traps

## Abstract

1. Insects are of crucial importance for terrestrial ecosystems but many populations decline rapidly. Conventional collecting methods are usually time-consuming, resulting in a low temporal, spatial and taxonomic resolution of data. Automated camera light traps (CLT) allow non-lethal monitoring of species-rich moths (Lepidoptera) and other nocturnal insects, but so far little is known about their performance compared to conventional collecting methods.
2. By observing the behaviour of moths in previous field work, we hypothesized that CLT perform well in moth groups in which species tend to sit down quietly after approaching the lamp (such as Geometridae) but worse in moth groups in which species are persistently active (such as Sphingidae).
3. We tested the performance of two CLTs, equipped with Sony alpha 7II (24 megapixel sensor) cameras that resulted in images with approx. 420 dpi resolution. The study was carried out in a forested area near Bielefeld in NW Germany for 196 nights in a row from April to September 2023, and photos were taken every two minutes during the night. All macromoths recognizable in the photos were identified and counted individually. We directly compared the data from the CLTs with moth samples obtained from conventional funnel light traps (FLTs) during 12 nights which were spread across the flight season.
4. The resulting 420 dpi images from the CLTs allowed reliable species identification of all observed macromoths with only few exceptions due to technical problems. In direct comparison during 12 nights, CLTs recorded 39 species exclusively, FLTs recorded 48 species exclusively, and 53 species were recorded by both methods equally. During the whole sampling period of 196 nights in a row, a total of 225 moth species were recorded by CLTs. CLTs tended to record Geometridae better than FLTs whereas Sphingidae tended to be undersampled by the new method. Familes differed in the length to which they remained on the screen of the CLTs.
5. Our study is the first to systematically compare the methods and it shows that CLTs perform overall very well. Results from CLTs differ to a certain extent from conventional trapping methods because they seem to perform worse in groups with highly active species and perform better in calmer groups like geometrid moths. CLTs are promising devices for insect monitoring since they deliver data with high resolution in time, space and taxonomy. The use of artificial intelligence (AI) for the analysis of images is intended as the next logic step.

## INTRODUCTION

Insects are of crucial importance for terrestrial ecosystems – as pollinators for wild plants as well as for many crops, as predators, decomposers or herbivores and generally as essential part of food webs (Noriega et al. 2018). However, there is growing evidence that many species are seriously threatened and in rapid decline (Wagner 2020, Warren et al. 2021). As an example, the well-known “Krefeld Study” revealed a decline in flying insect biomass of 76 % over 27 years (Hallmann et al., 2017). This decline was recorded within German nature conservation areas and could not be explained by habitat type, weather conditions or land use changes. Since then, insect declines have been reported worldwide – most evidence comes from western and northern Europe, but also from Asia, North America, the Arctic, Neotropics, and elsewhere (Wagner 2020, Conrad et al. 2006). Population declines in insects are probably driven by many factors, including climate change, habitat loss (or degradation), synthetic pesticides and fertilizers as well as light pollution (Sánchez-Bayo & Wyckhuis 2019, Van Sway et al., 2006, Brehm et al. 2021).

The vital importance of insects in ecosystems and their substantial decline call for a profound understanding of insect population dynamics. However, large-scale, continuous and species level data sets on insect populations are largely missing (Bjerge et al. 2021, Conrad et al. 2006). An important reason is that conventional methods are usually (very) time-consuming, resulting in either high personnel costs or low resolution in terms of temporal, spatial and taxonomic accuracy (Bjerge et al. 2021). Therefore, new methods are needed that are fast and efficient, yet reliable and with a high taxonomic resolution, and ideally without negative effects such as the killing of individual insects.

Among insects, butterflies are probably the best-known insect group and monitoring programs have been established across many countries, especially in Europe (Van Swaay et al. 2008, Dennis et al. 2017). However, butterflies are a relatively species-poor insect group in many regions, and monitoring can usually only be carried out with a relatively high workload because they must be individually observed and recorded. Butterflies make up around ten percent of all Lepidoptera (van Nieukerken et al. 2011), while the remaining majority (usually referred to as “moths”) comprises usually less conspicuous species that are mainly active during the night. The so called macromoths alone – representing the fraction of larger moth species – are represented with around 1160 species in Germany (Steiner et al. 2014).

Moths appear suitable for monitoring for three reasons. First, they are one of the very few large and species-rich insect groups that can be identified comparatively well, based on macro-optical characteristics such as wing markings (e.g. Steiner et al. 2014) – thus, they can well be detected with a camera sensor. In contrast, the identification of many other insect species requires specialized knowledge and a close-up look of the insect itself, the preparation of their genitalia and / or molecular methods. Second, moths represent a significant proportion of the native insect fauna (Steiner et al., 2014, Völkl et al., 2004). In central Europe, there is a comparatively detailed knowledge of their ecology, meaning they can serve as indicators and are thus of great value for practical nature conservation. Furthermore, since moths are diverse in their ecological requirements and deploy a wide range of habitats and host plants they can be considered as representatives of other insects or entire bioscenoses (Steiner et al., 2014, Van Swaay et al., 2006). The third reason why moths are suitable for monitoring is that they can be easily and effectively captured using UV radiation (Brehm, 2017). Nocturnal insects use natural light gradients to maintain proper flight attitude by tilting their dorsum towards the brighter visual hemisphere, which is under natural conditions always the sky (Fabian et al. 2024). However, in the vicinity of artificial light sources this behaviour can result in the moths no longer being able to escape. UV radiation is known to strongly attract a wide variety of insects and has become particularly well established for recording nocturnal moths (Jonason et al., 2014, Brehm et al., 2021). So far, moths were either collected manually (e.g. UV lamp and sheet) or semi-automatically (e.g. using funnel traps) (Brehm & Axmacher, 2006). In the latter case, a UV lamp is installed above a funnel and moths are captured in a net or bucket placed below (Singh et al., 2022), and traps must be controlled after every night of collection.

So far, camera traps have mainly been used to track vertebrates and mammals, particularly those with a rather discrete way of life (Harley & Eyre, 2024, Russo et al., 2023). Also for insects, automated monitoring methods using visual systems have been increasingly used in recent years, but often limited to study pest species (Preti et al., 2021). Various papers have described automatic traps for pest monitoring, usually consisting of a trap structure (e.g. with a bait), and a box including the camera, electronics, data storage and a battery or external power supply (Preti et al., 2021). For setting up a camera trap for nocturnal insects, we have been continuously tested a system in a dry grassland site in Jena-Ziegenhain (Germany) since 2020 (Korsch et al., 2023; GB, unpublished data). The device is here referred to as camera light trap (CLT), very similar in its basic principle as shown in Bjerge et al. (2021). It consists of a tripod with a camera on one side and a UV lamp mounted over a screen (where moths are photographed) on the other side (see methods). We counted and identified species manually for this study but are well aware that these tasks will be taken over in future studies by deep learning algorithms (Bjerge et al., 2021, Korsch et al., 2023). Our work aims to generally contribute to better understanding of moth monitoring with CLTs. Previous studies were limited by a relatively low effective image resolution with the effect that only large and conspicuous species could be identified (Bjerge et al. 2021, Möglich et al. 2023). We were using a higher resolution of about 420 dpi, to test if all macromoth species can in this way reliably be identified. Moreover, our two main hypotheses are based on observations made during field work in previous years but which have not yet been quantified:

1. We expect that certain Lepidopteran taxa can be captured as well with CLTs as with conventional FLTs, because they quickly calm down on the screen and also remain quiet for a certain time. This includes in particular geometrid moths (Geometridae). It might even be possible that these taxa can be better assessed because they are relatively small and lightweight, and are thus not necessarily captured with a FLT because they remain sitting on the outer parts of the trap and leave in the early morning hours as described by Brehm & Axmacher (2006).
2. On the other hand, we expect that certain other moth groups are less well recorded by CLTs than by FLTs. We expected this in species that are known to be highly active once they have started flying. For example, hawkmoths require high body temperatures (Heinrich, 1993) and are often continuously active after they have come into the vicinity of a UV lamp, making it relatively unlikely that they will be detected by CLTs, but more likely to be captured by FLTs.

## MATERIALS AND METHODS

### Study site and habitats

The study area is located in NW Germany (North Rhine-Westphalia), near Schloss Holte-Stukenbrock (ca. 51.9° N, 8.7° E, 130–140 m asl.). The larger region is dominated by intensively used agricultural fields and the area is part of the old cultural landscape Senne, which is the largest nutrient-poor sandy landscape in the larger region. We chose two monitoring sites on the land of farmer Gerhard Brechmann, which is part of the nature conservation area Wehrbachtal. The area of 13 ha was designated as conserved area in 1990 and consists of a mixture of forested and open habitats. Riparian alder-ash forests, old oak-beech stands, wet meadows and dry grasslands should be mentioned here as particularly valuable structures (Brechmann 2023).

Each one CLT and FLT were set up in two sites in different habitats at a distance of about 200 m (detailed GPS data of sites in supplementary material Table S1). The open habitat was a small meadow in the direct vicinity of the main farm building (Fig. 1). The meadow is surrounded by forest dominated either by beech, oak and hornbeam. Shrub vegetation is dominated by hazel, hawthorn, bird cherry, blackthorn, elderberry and honeysuckle. The meadow itself is largely dominated by various grasses, however, also other typical caterpillar host plants like cuckoo flower, cleavers, common ragwort, plantain species and various dock species are found. The forest habitat site was located in pine-dominated forest, with also occurring birch, rowan and alder buckthorn. The herbaceous layer comprised species such as may lily and blueberry species indicating the nutrient-poor and sandy soils in the Senne.

**Fig. 1.**
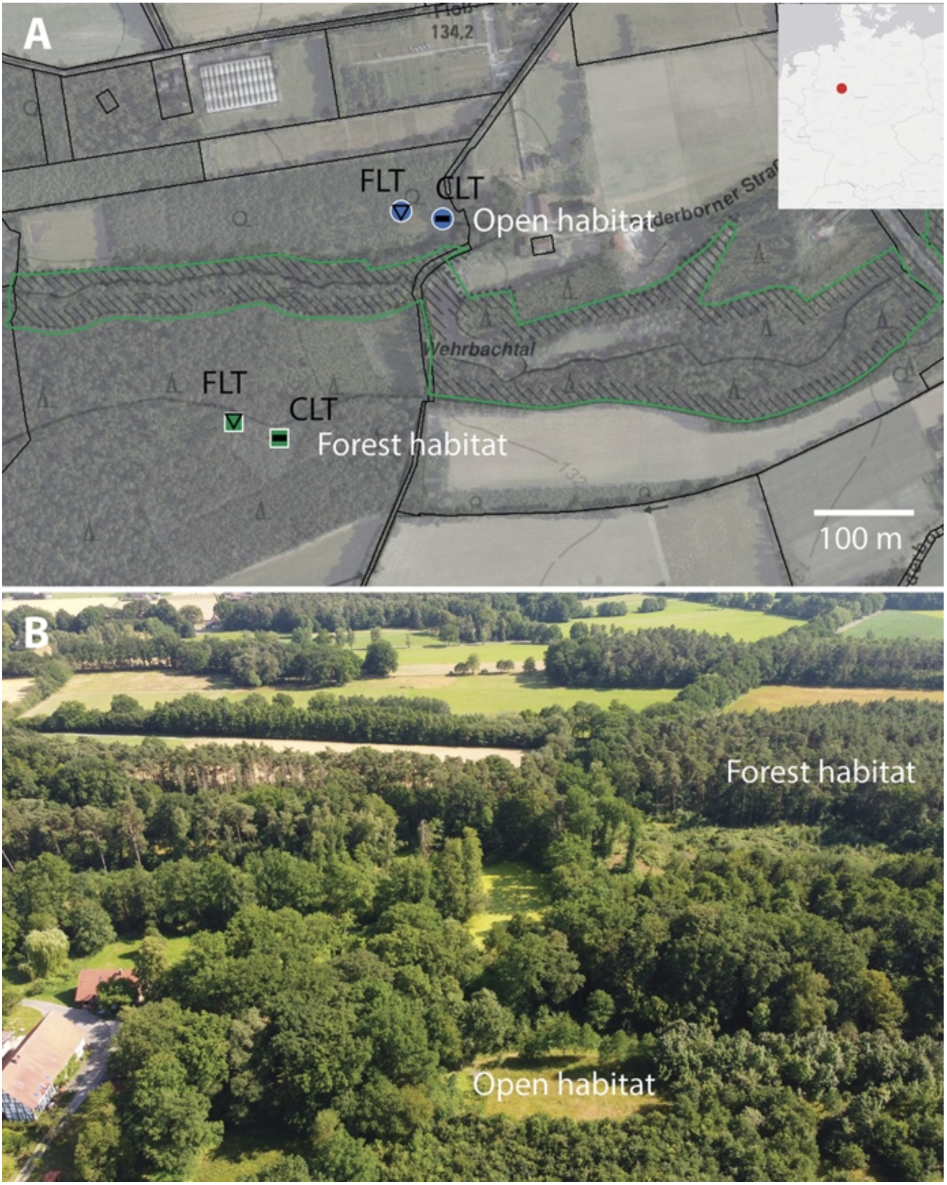
**A** Map of study area in NW Germany. FLT = Funnel light trap, CLT = camera light trap. Blue symbols: open habitat, green symbols: forest habitat. Source: GEObasis.nrw. **B** Aerial photograph of study area, taken with a drone facing south (8th July 2023).

### Camera light traps and funnel light traps

Both trap types were operated using the same UV lamp (LepiLED Mini switch from Insects & Light, Jena, Germany). The FLTs were powered by 26 Ah power bank batteries during night. Due to the direct vicinity to the main farm building, the open habitat site could be supplied with mains power. In the forest, die CLT was powered by a Varta Silver Dynamic car battery (12 V, 100 Ah), which was changed and recharged regularly.

The CLT was constructed with 15 mm aluminum profiles mounted on a wooden frame. The UV lamp was mounted above the screen, made of a 10 mm polyethylene foam glued to an aluminium plate. We used a full frame Sony alpha 7 II mirrorless camera (24 megapixel sensor) with a Sony lens (f = 50 mm SEL50M28). Data were locally stored on 512 GB San Disk Ultra SD cards (exchanged together with the battery every ca. ten days). The camera was set to a distance of approx. 50 cm so that only the screen and a narrow surrounding edge were photographed. With an image width of 6000 pixels (height 4000 pixels) and a screen width of 36 cm, this results in a resolution of around 165 pixels per cm (= ca. 420 dpi). Photos were taken every two minutes between dusk and dawn. We used a Neewer TT 560 flash for lighting. The main settings of the camera were ISO 200, with manual focus, f/11 in A (automatic) mode, taking fine jpg images. We chose manual focus after testing different focus settings,but noted that it was necessary to block the contact between lens and camera because otherwise the camera slightly shifted in focus when it was “wakened” in the evening from resting phase, resulting in blurred images. Shutter releases and UV lamp were controlled with a Siemens logo device according to fixed preset times. The cameras were switched on at sunset, while UV lamps were switched on half an hour before, and both devices were switched off at sunrise. Timer, cables and batteries were stored in a waterproof box unterneath the CLT. A roof of transparent polyethylene was built in order to ensure protection of the equipment against rain, hail and falling branches. Camera, lens, flash and cable connections were additionally wrapped into foil for further protection. The mandatory maintenance of the CLTs was limited to only a few tasks: In intervals of approx. ten days, SD cards had to be exchanged, car battery had to be recharged and timers were regularly adjusted.

The FLT (Insects & Light, Jena, Germany) were attached to larger tree branches (Fig. 2B) and had the same design as described by Singh et al. (2022). They consist of a roof (diameter 22 cm) with a carabiner for suspension, three vertical vanes that connect roof and funnel and the funnel itself (lower diameter 80 mm). Moths and other insects were captured in a black mesh bag with a capacity of 60 l hooked into the margins of the funnel. Egg cartons were placed in the bag to provide sufficient space for the moths to sit and hide.

**Fig. 2.**
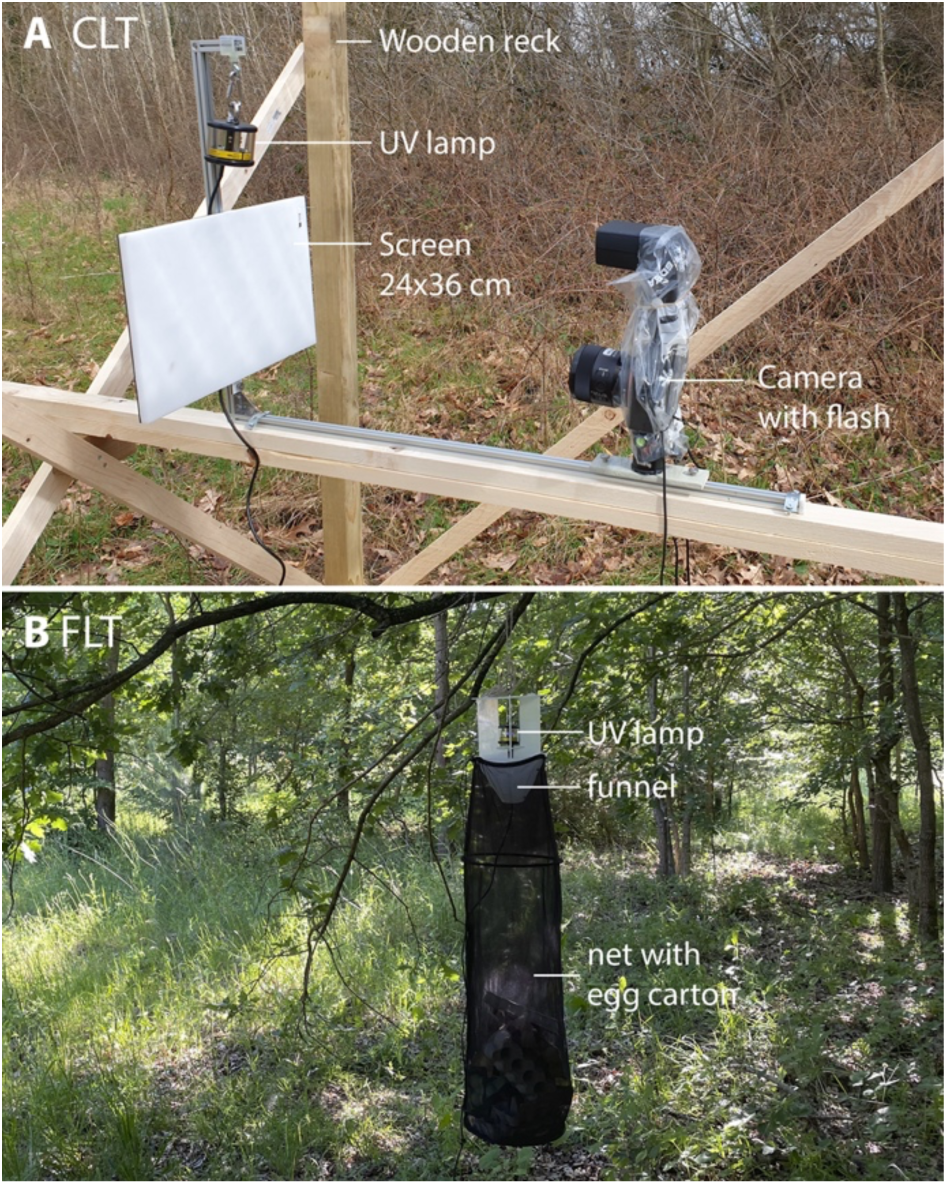
**A** CLT = Camera light trap, built on a temporary wooden rack and protected from rain with a transparent roof (not shown). **B** FLT = Funnel light trap, with 60 l black net, filled with egg cartons.

Collection with FLTs was conducted twice per month from April until September (12 times in total), in a timespan of ± five days around new moon. The specific date for trapping was matched with weather conditions so that light trapping was preferably performed in mild and cloudy nights, with only little rain or wind (supplementary material Table S2). FLTs were placed in the field in the late afternoon to evening with a timer set, but not before 4 pm. In the following morning, the FLTs were emptied between 7 and 8 a.m. Captured specimens were identified and photographed directly in the field and, in most cases, released. In both habitats, FLTs were positioned approx. 20 m from the CLTs (Fig. 1A) to prevent interaction and interference between the devices. It was also ensured that the two trap types were still in similar habitat with comparable vegetation, despite the distance. FLTs were hung in such a way that the UV lamp had the same height as in CLTs.

### Image processing and species identification

Image data were copied every ca. ten days onto a computer hard disk, checked and reviewed manually if insects were present on each photo. For uncertain cases we used the identification app Obsidentify (Stichting Observation International, 2023). Obsidentify suggestions were then critically checked for plausibility with the German website Lepiforum (Rennwald & Rodeland, 2023) as well as with a comprehensive field guide (Steiner et al. 2014). Species that could not be determined with certainty, e.g. due to the specific situation (only partially visible or blurred due to rapid movement, etc.) or in principle (species that require the use of genital morphological features for identification) were marked accordingly and considered separately.

A few species complexes remained unresolved for statistical analyses. Some individuals in FLT samples could be identified at species level, whereas some individuals CLTs were only identified at the level of a species complex. For consistency, we subsequently used the same species complexes from CLTs also in FLT samples. Specimens that could only be determined to genus level as well as microlepidopterans were excluded from the analyses. For CLT data, we counted observations as a substitute for abundance. An observation is here defined as a sequence of the same species photographed in succession. The time between two photos was usually two minutes, but in some cases up to three minutes. Thus, we set the time difference threshold for one appearace between two consecutive observations for each species to be smaller than four minutes.

We declared photographs as defective if this led to depicted moths not being able to be further identified. For example, there were cases in which several photos in a row were blurred, but a single or more subsequent photographs showing the same moth specimen in focus. In such cases, it was possible to include data even from blurred images. In other cases, moths could not be identified, and observations had to be excluded from analysis. In total, this led to approx. 1,000 excluded observations.

### Statistical analyses

All statistical analyses were performed using R (version 4.3.2, R Core Team, 2021) and RStudio (version 2024.04.1+748, RStudio Team 2024). We applied the “vegan” package (Okansen et al, 2020) for analyses of the diversity. For estimations of species numbers, we calculated the Chao estimator (Chao, 1987), Jackknife 1-index (Zahl, 1977) and the bootstrap estimator (Burnham and Overton, 1979). We calculated species accumulation curves for a direct comparison between simultaneous samples of both CLT and FLT as well as for all CLT samples throughout the whole season by using the “iNEXT” package (Hsieh et al, 2016). We checked for shared species between the trap types and visualized the results with a Venn Diagramm using “ggVennDiagram” (Cao et. al, 2021). We ordinated the species composition of samples collected simultaneously in both habitat types by both trap types with a non-metric multidimensional scaling (NMDS). We further checked for differences in community composition with an analysis of similarities (ANOSIM) with 9999 permutattions between both trap types. We conducted an analysis of indicator species with the “indicspecies” package (de Cáceres and Legendre, 2009) to check for species that are associated with one of the trap types. For comparisons of observed individual numbers, we generated a pairwise boxplot for all families of Macrolepidoptera. We visualized the accumulated observation time for the ten longest observed species and checked for the top three most observed species. Additionally, we checked for differences in durations of all appearences amongst all indivuduals of all families. We generated a boxplot representing all individuals per family and used a pairwise Wilcoxon test.

## RESULTS

### Moth diversity recorded by CLTs and FLTs

The CLTs in the forest and open habitats were operated for a total of 196 nights between April and September 2023. FLTs were operated 12 times, covering all new moon phases in that period. With only few interruptions, the two CLTs quite reliably produced photographs during the entire sampling period, resulting in a total of approx. 120,000 photographs. Over the course of the season, around 1,700 photographs at both locations (1.4 % of all photographs) could not be analyzed due to technical problems concerning the focus (blurry images) and around 7,200 photographs were lost due to defective triggering (6.0 %).

Macromoths could generally be well identified with the given resolution of approx. 420 dpi (Fig. 3). In a first step, 252 operational taxonomic units of Lepidoptera were recorded (spreadsheed in supplementary material). Ten Microlepidoptera species and four genera without species identification were not included in the subsequent analyses, i.e. the final dataset comprised 238 species / operational taxonomic units, including twelve unresolved species complexes (spreadsheed in supplementary material). Numbers of individuals and species, broken down by moth family and habitat are provided in Table S3 in supplementary material.

**Fig. 3.**
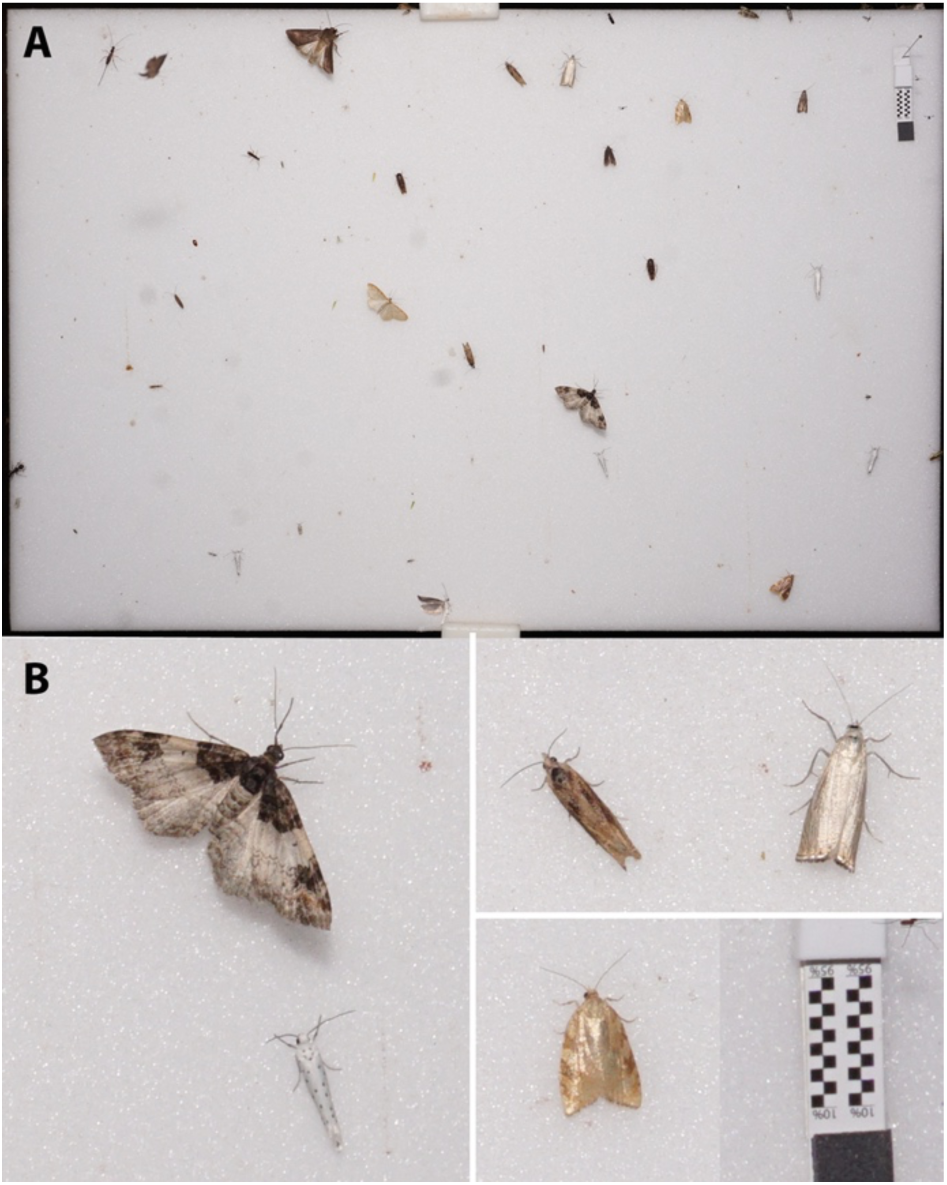
Example of a photograph taken with a camera light trap (CLT) at 420 dpi. **A.** Overview of complete screen (24 x 36 cm). **B.** Magnified details. Scale bar 10 mm. Further examples are provided in supplementary material Fig. S1.

In direct comparison, moth richness and diversity recorded by FLTs was slightly larger than recorded by CLTs (Fig. 4A, supplementary table S4). Over the entire season however, CLTs recorded a far higher species richness than both trap types during 12 selected nights (Fig. 4B). Moth diversity showed a peak from beginning of June to mid-July, which coincides with the peak season for most species, although a brief decline in numbers was recorded at the beginning of July (supplementary material Fig. S2). It also appears that in the first half of the season, the forest site revealed more observations, while in the second half the open habitat became more dominant. A comparison of the performance of the two trap types at family level revealed similar observed species numbers in both methods (Fig. S3 in supplementary material).

**Fig. 4.**
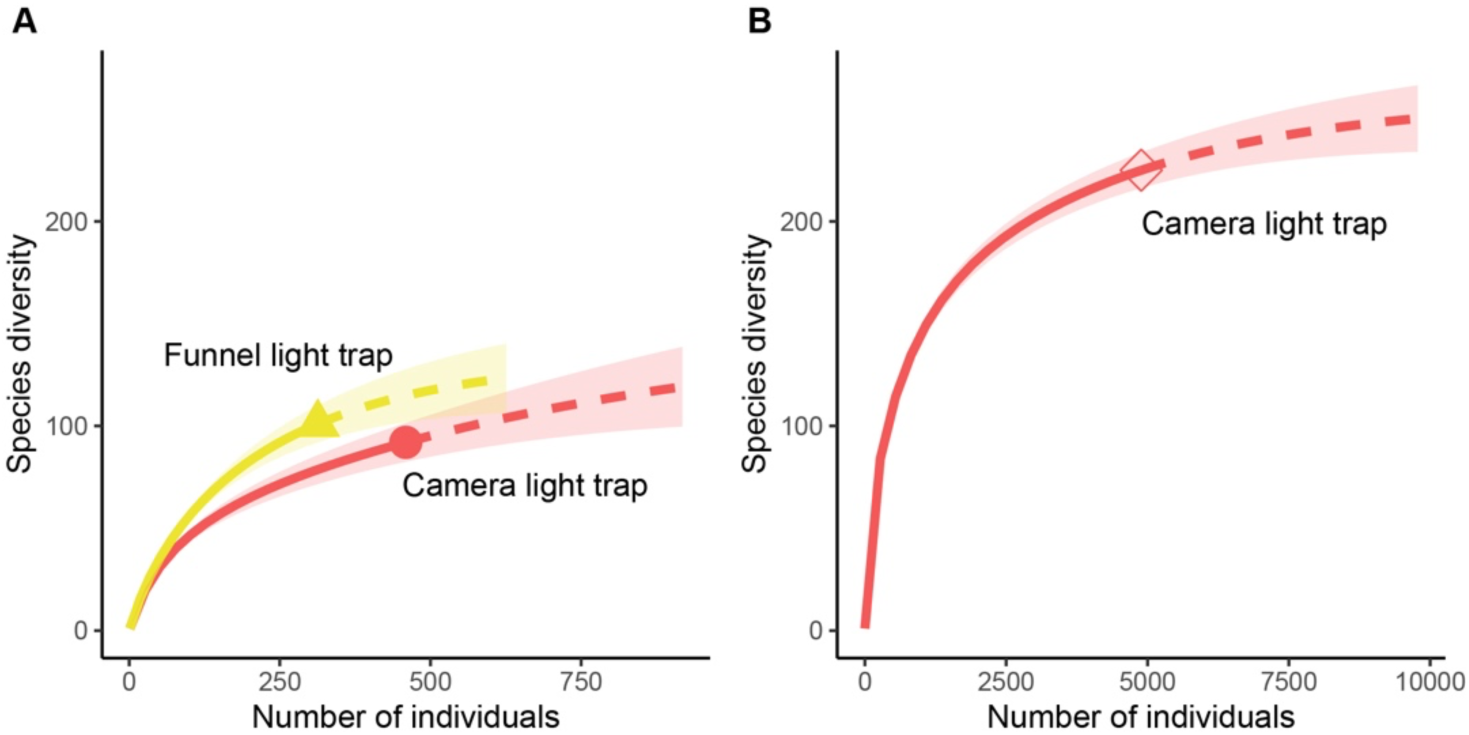
**A.** Moth species diversity comparison of 12 trapping nights in which both trap types (FLT and CLT) were operated. **B.** Moth species diversity of 196 nights in which the CLTs were operated.

140 species of moths were recorded in total in direct comparison during 12 nights. CLTs recorded 39 species exclusively, FLTs recorded 48 species exclusively, and 53 of the species were shared (Fig. 5A). In 196 nights in a row, a total of 238 moth species were recorded. 137 moth species were recorded exclusively with CLTs as compared to 13 species recorded exclusively by FLTs, and 88 species were shared (Fig. 5B).

**Fig. 5.**
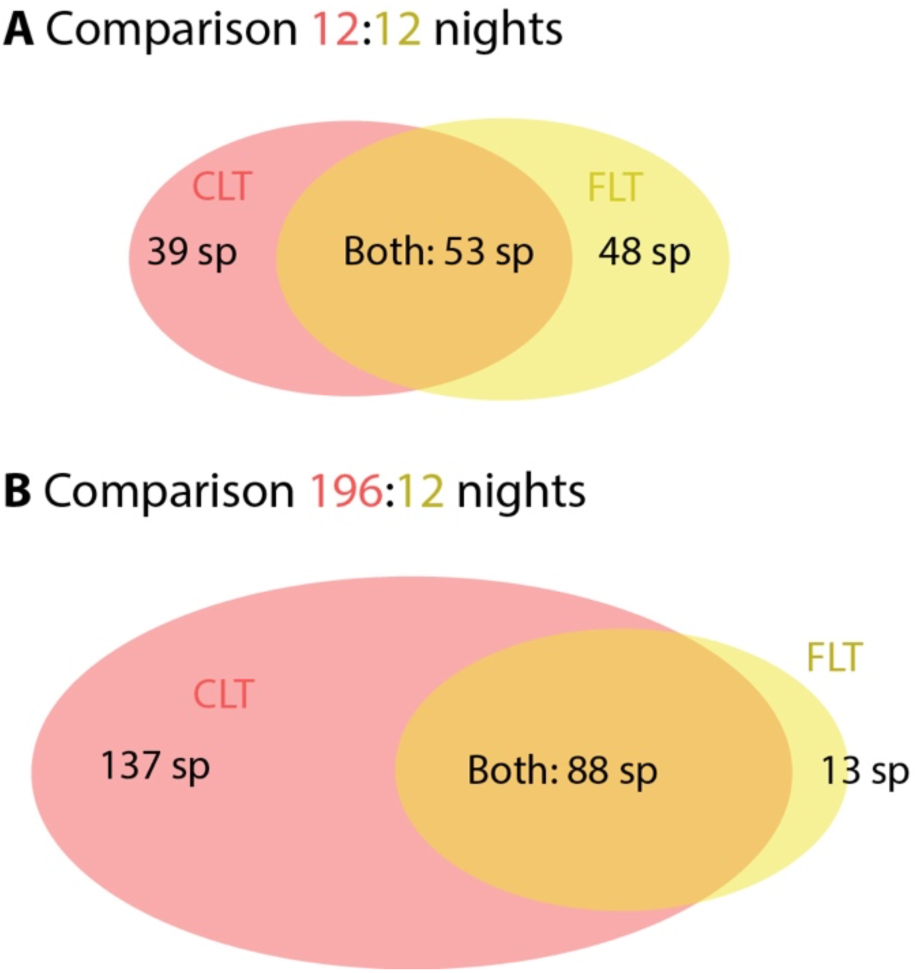
Venn diagrams showing the number of species detected by CLTs (light red) and FLTs (yellow) each method as well as shared species. **A.** comparison of 12 trapping nights in which both trap types were operated. **B.** Comparison of 196 nights in which the CLTs were operated with 12 trapping nights in which FLTs were operated.

### Moth communities recorded by CLTs and FLTs

In direct comparison (12 vs. 12 nights, samples of the two habitats combined), moth communities recorded with CLTs and FLTs were largely overlapping (Fig. 6). An Anosim between the communities captured by both trap types did not show differences in community composition (p = 0.073; R = 0.099). The ordination revealed a relatively clear timely pattern: Early year moth communities are more likely to be found in the left and lower part of the ordination, whereas summer and autumn communities tend to be found in the upper and middle part of the ordination.

**Fig. 6.**
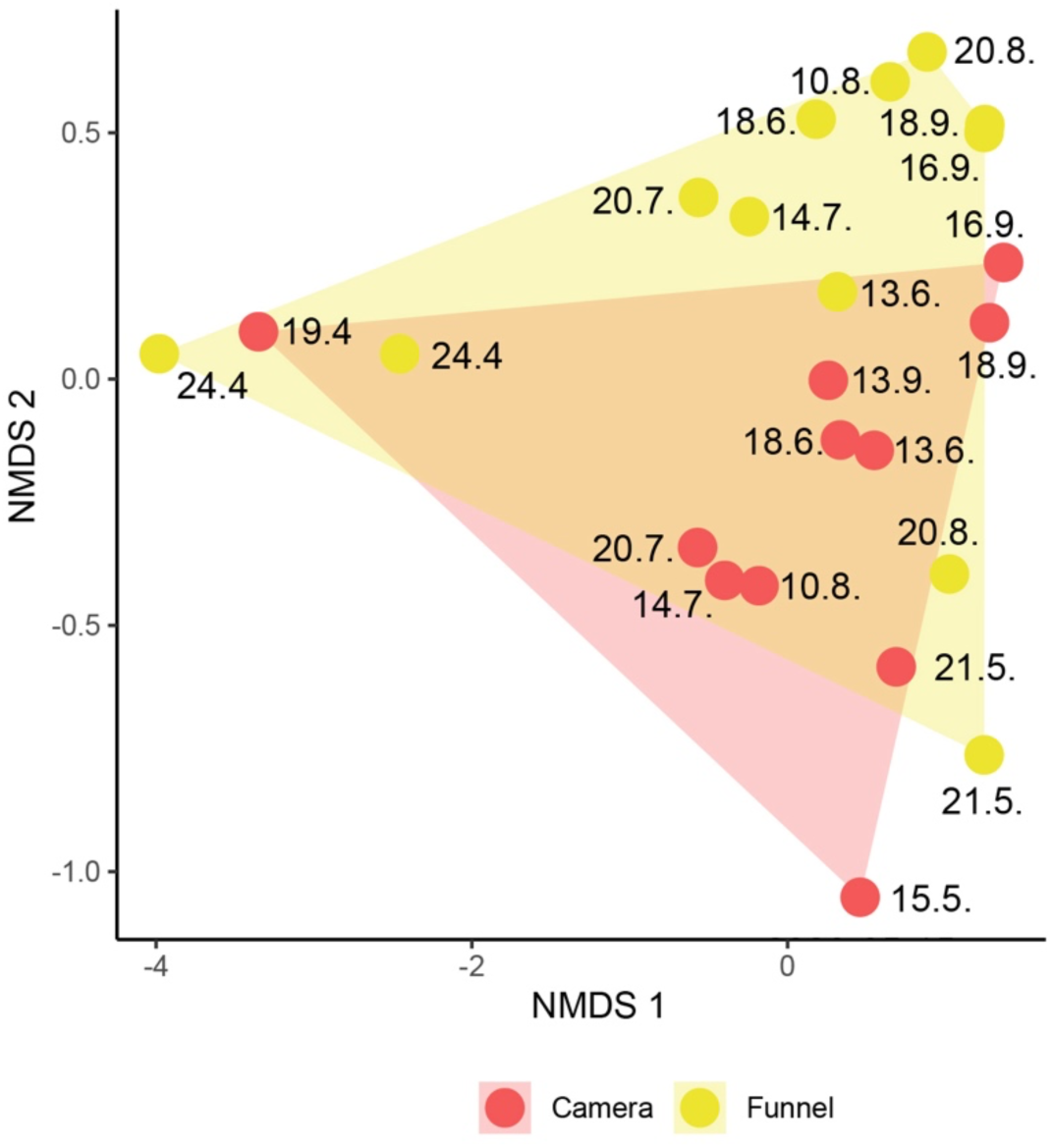
Non-metric multidimensional scaling (NMDS) of 24 species assemblages, gained from two trapping methods (CLTs and FLTs) in two habitats (combined) during 12 nights. All assemblages have a date (all in 2023).

We found six indicator species for CLTs (all Geometridae) and one species for FLT, a hawkmoth species (Table 1). At the level of families, Geometridae were more often observed in CLTs than in FLTs (supplementary material Fig. S3). Differences at family level were not significant, but hawkmoths (i.e the pine moth *Sphinx pinastri*) were observed in five out of twelve times in FLTs but were not detected by CLTs at all.

**Table 1.**
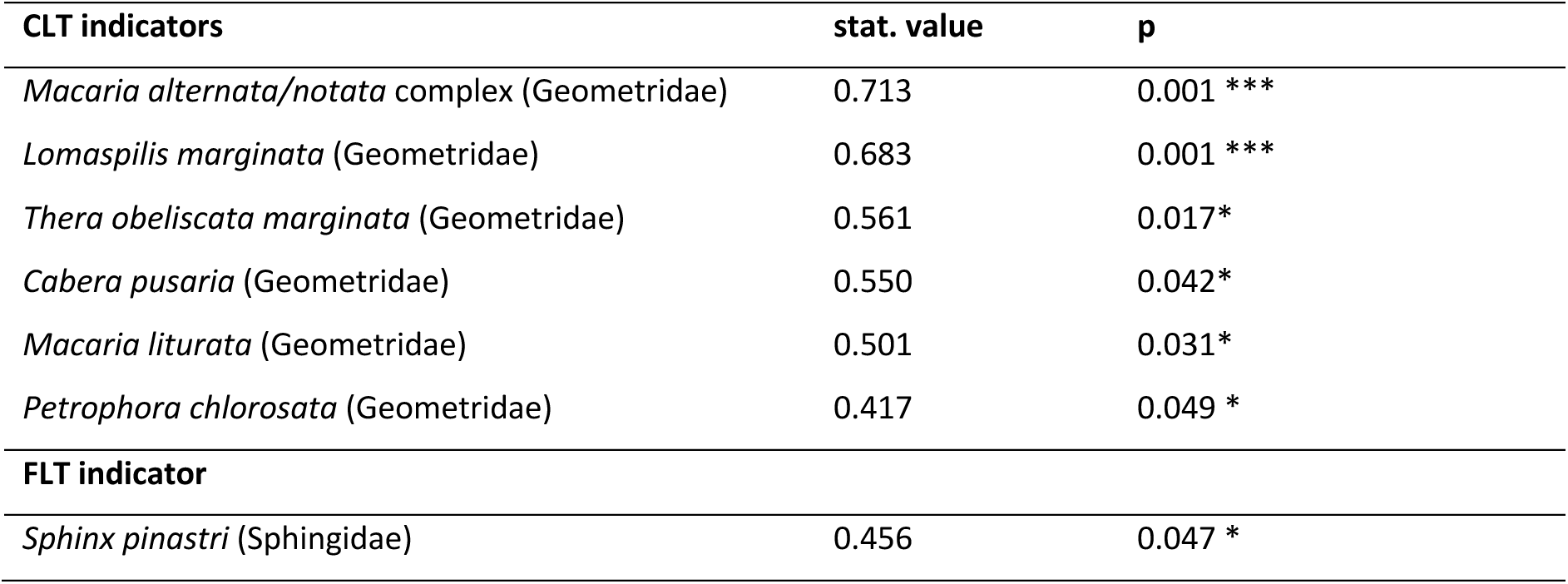
Indicator moth species of camera light traps (CLTs) and funnel light traps (FLTs).

| CLT indicators | stat. value | p |
| --- | --- | --- |
| *Macaria alternata/notata* complex (Geometridae) | 0.713 | 0.001 *** |
| *Lomaspilis marginata* (Geometridae) | 0.683 | 0.001 *** |
| *Thera obeliscata marginata* (Geometridae) | 0.561 | 0.017* |
| *Cabera pusaria* (Geometridae) | 0.550 | 0.042* |
| *Macaria liturata* (Geometridae) | 0.501 | 0.031* |
| *Petrophora chlorosata* (Geometridae) | 0.417 | 0.049 * |
| **FLT indicator** |  |  |
| *Sphinx pinastri* (Sphingidae) | 0.456 | 0.047 * |

### Duration of observation sequences in CLTs

The three species with the highest number of observation sequences are *Lomaspilis marginata* (Geometridae: 474), followed by *Eilema sororcula* (Erebidae: 213) and *Deltote pygarga* (Noctuidae: 201). However, these are not necessarily the species that spent most total time on the screen, as these three species are *Lymantria dispar* (Erebidae), followed by *Pennithera firmata* (Geometridae) and *Campaea margaritaria* (Geometridae) (Fig. S4 in supplementary material). Some species were primarily observed in the forest, e.g. *P. firmata*, or in open habitat, like *C. coryli*, while other species such as *L. marginata* and *Pheosia gnoma* were similarly long visible in both habitats (Fig. S4 in supplementary material). When considering the longest median observation sequence durations, *Chloroclysta siterata* (Geometridae, 630 minutes, N = 3), *Cerapteryx graminis* (Noctuidae, 464 minutes, N = 1) and *Photedes fluxa* (Noctuidae, 412 minutes, N = 2) lead the ranking.

The family with the most observed sequences is Erebidae, followed by Geometridae (Fig. 7). Noctuidae and Drepanidae were also frequently observed, but in a different order of magnitude than the former. Members of Notodontidae had by far the longest average and median residence times. They are also the only family without outliers, which are present in all other groups (Fig. 7). Lasiocampidae consistently show the lowest residence time. Differences between Notodontidae and most other families are significant (except for Limacodidae and Nolidae, Wilcoxon-Test). There is also a statistically significant difference between the residence times of Noctuidae and Drepanidae (Fig. 7). Boxes of almost all families appear similarly “compressed” at low levels, indicating that most of the associated observation sequences were short. The median duration of observation sequences is in most cases 2 or close to 2 (supplementary material Table S6).

**Fig. 7.**
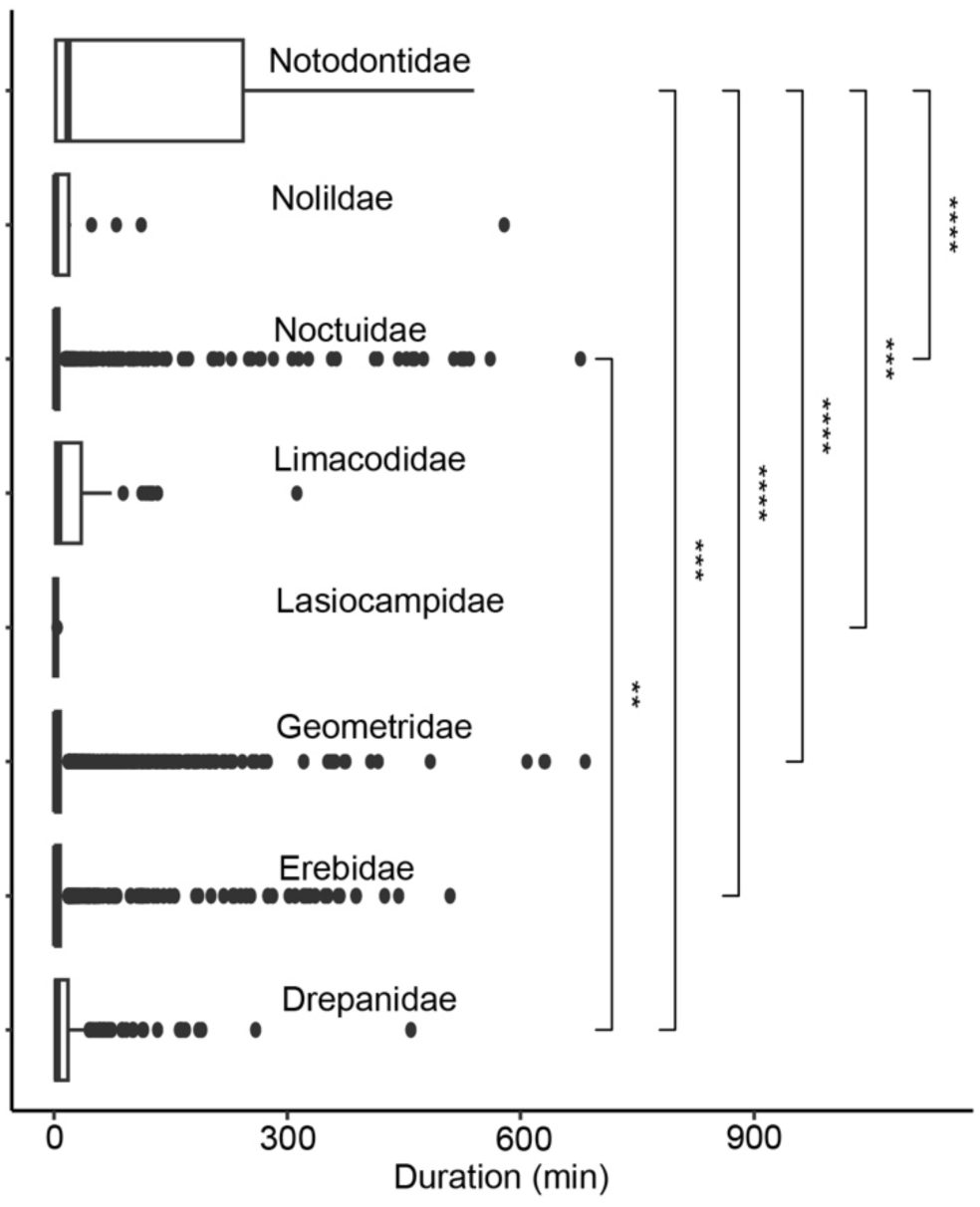
Boxplot showing the observation sequence duration by family. Plotted are the upper and lower quartiles (boxes), medians (horizontal lines), standard deviation (error bars) and outliers. Square brackets indicate significantly different pairs (Wilcoxon-Test, p < 0.05).

## DISCUSSION

### General performance of 24 megapixel CLTs

Our CLTs generally performed well, despite some technical problems (discussed below). In particular, it was possible to generally identify macromoth species, including also many of the small species such as *Eupithecia* spp. and *Idaea* spp. Twelve species complexes could not be resolved because the species can generally not be distinguished reliably by a photo or the species were not in a good condition (supplementary material Table S5). The CLTs of our study used cameras with 24 megapixel sensors and decent lenses, and thus a potentially high effective resolution. Previous studies by Bjerge et al. (2021) and Möglich et al. (2023) were using webcam technology with far lower effective resolution (< 8 megapixel sensors) combined with a larger screen. This lead to the result that only larger species could be identified with reasonable certainty in these studies. A higher resolution is obviously advantageous, as it allows the identification of the large majority of macromoths – and presumably also the identification of many microlepidopterans and other insects in the future (Fig. 3). The most important trade-off is between the largest possible sensor size and the largest possible screen size on the one hand and the lowest possible price for the system and the lowest possible storage requirements on the other. There is no universal answer to this question, but we believe it is fundamentally important that certain target groups (in our example, macromoths) can be captured reliably. We have achieved very good results with a resolution of approx. 420 dpi with the selected size of image sensor and screen. Future studies should determine more precisely whether similar results could be achieved with a slightly smaller sensor (as found in many industrial cameras), for example.

Disadvantages in the use of mirrorless cameras are that they are expensive and not built for permanent use in outdoor conditions. As these cameras can be used for many purposes, risk of theft is also greater than with devices that can only be used by specialists. In the medium term, the use of high resolution webcams or industrial cameras appears therefore a preferred choice, as these are cheaper and also take up less space to be housed in a casing (see discussion below).

CLTs allow very detailed insights into the phenology of insect species that can hardly be deteced by conventional methods. For example, the gypsy moth *Lymantria dispar*, a potential pest species, was the most frequently observed species. It was detected on more than 3000 photographs, and was the species with the longest total visibility time. *L. dispar* dispersed only recently into the region where it was observed in 2010 for the first time (Pähler & Dudler 2013, Schulze et al. 2022), and it now seems to be one of the commonest species.

### Performance of CLTs versus FLTs

In direct comparison of 12 collecting nights, CLTs delivered similar results to FLTs – in detail, however, we were able to quantify in our study what we had already observed in previous years. We only found trends, but some of these could be significant in future studies with larger samples. It seems likely that Geometridae can on average be recorded even better overall with the CLT method than in conventional traps because all six indicator species for CLTs were geometrids. On the other hand, it was shown that hawkmoths can be detected less well by CLTs than by funnel traps. However, our data are based on observations of only one species, namely *Sphinx pinastri*. Here, too, we hope that future studies in more species rich regions will allow clearer conclusions to be drawn. Our data show that although the results are generally comparable between the methods, they are not identical. When analysing further data from CLTs, it should therefore be taken into account that certain moth groups cannot be recorded well using this method. It remains to be seen whether this applies primarily to Sphingidae or whether other groups are also affected – such as Lasiocampidae and other families with highly active species. It is a simple fact that there is no survey method for any major animal group that provides unbiased data on species abundance – attraction by UV radiation is probably one of the most universal collection methods in animal ecology. Given the overall similar number of species detected with CLTs and FLTs, we see no fundamental problems in using CLTs as a monitoring method.

CLTs offer an efficient alternative to much more laborious and time-consuming conventional methods. Once the CLTs and the related technology have been successfully set up in the field, they work independently and require a comparatively low level of supervision. Conventional methods such as manual light trapping with sheets require a person to spend all or part of the night in the field (Brehm & Axmacher 2006). Semi-automatic traps, such as FLTs, require far less effort, but must be checked at least once a day, preferably in the early morning. CLTs are easily able to produce large, continuous data sets – as in the present study in 196 consecutive nights. Such long-term effort has only been achieved in relatively few manually monitoring schemes such as the Rothamstead light trap network, involving a high number of volunteers. Moreover, CLTs make it possible to precisely record the occurrence of insects over the course of a night whereas FLTs usually represent the sum of all individuals and species collected during a night.

Ethical aspects play an increasing role also for invertebrates (Fischer & Larson, 2019), and they are generally more in favor of CLTs than FLTs and comparable methods. All light trapping methods basically mean a disturbance of nocturnal insects and a potential loss of their fitness. This can be well justified by the acquisition of data on the occurrence and abundance of insects, as without such data important foundations for decisions in nature conservation are simply missing. Apart from that, placing any artificial light source in the landscape has a similar effect on insects – but without any positive aspect for them. Although both presented methods avoid killing insects, moths (especially very active ones) are more likely to damage themselves in a net or bucket than on a CLT.

Can CLTs be as precise as conventional methods? In the future, this will largely depend on how good AI models are (see below). The CLT method reaches its limits in cases where characteristics cannot be assessed by means of the appearance of individuals. In some cases, therefore, the species level is not reached, but the level of species complexes (e.g. in *Mesapamea secalis* / *secalella*). However, this is only a problem in relatively few cases in the Central European fauna. Species groups can in principle be treated statistically in the same way as species. If a more precise breakdown of such complexes is required, other methods must be used, such as dissecting the genitalia of the moths or analysing gene sequences.

As all camera data was still analysed manually in our study, the amount of work involved was high. Of course, this would have been orders of magnitude greater if a large number of CLTs had been in use. The use of AI is therefore the next logical step, as a large amount of comparable standardized data is available here, which is generally very suitable for evaluation with AI (see below; Bjerge et al. 2021, Möglich et al. 2023).

### Technical aspects and future perspectives of CLTs

Although we were overall satisfied with the performance of the CLTs, there were some technical problems in our setup that should be avoided in future designs. The control of the device was overall rather simple and required multiple adjustments due to the changed day length. The use of flash lights produced good image results at short shutter speed, but we have already had problems in several cases because the flash units did not work. Frosty nights in spring can be a general problem for the technology used as it is usually not designed for a temperature range below 0° C. Furthermore, there will always be a certain percentage of moth photographs that cannot be interpreted, e.g. because the moth is in motion or can only be partially seen.

With new technical approaches most of the problems mentioned can probably be minimized. The new and further developed LEPMON CLTs have the same fundamental design as our CLTs (Fig. 8). The main phase of the project started in December 2024 and will involve installing more than 100 CLTs across Germany (DB, GB, PB, unpublished data). These devices are controlled using a Raspberry Pi, which allows many functions to be controlled far more precisely than with the CLTs we have used in our study. This includes, among other things, the twilight-dependent control of the light and the camera, but also the integration of light and temperature sensors and the naming of the image files according to a fixed scheme. A cheaper and more compact industrial camera is used so that all the electronics can be housed in a waterproof housing (Fig. 8). The housing and screen are firmly mounted on an aluminum frame so that the system can be easily set up in the field. A detailed description of the LEMPON CLT is in preparation (DB, GB, unpublished data).

**Fig. 8.**
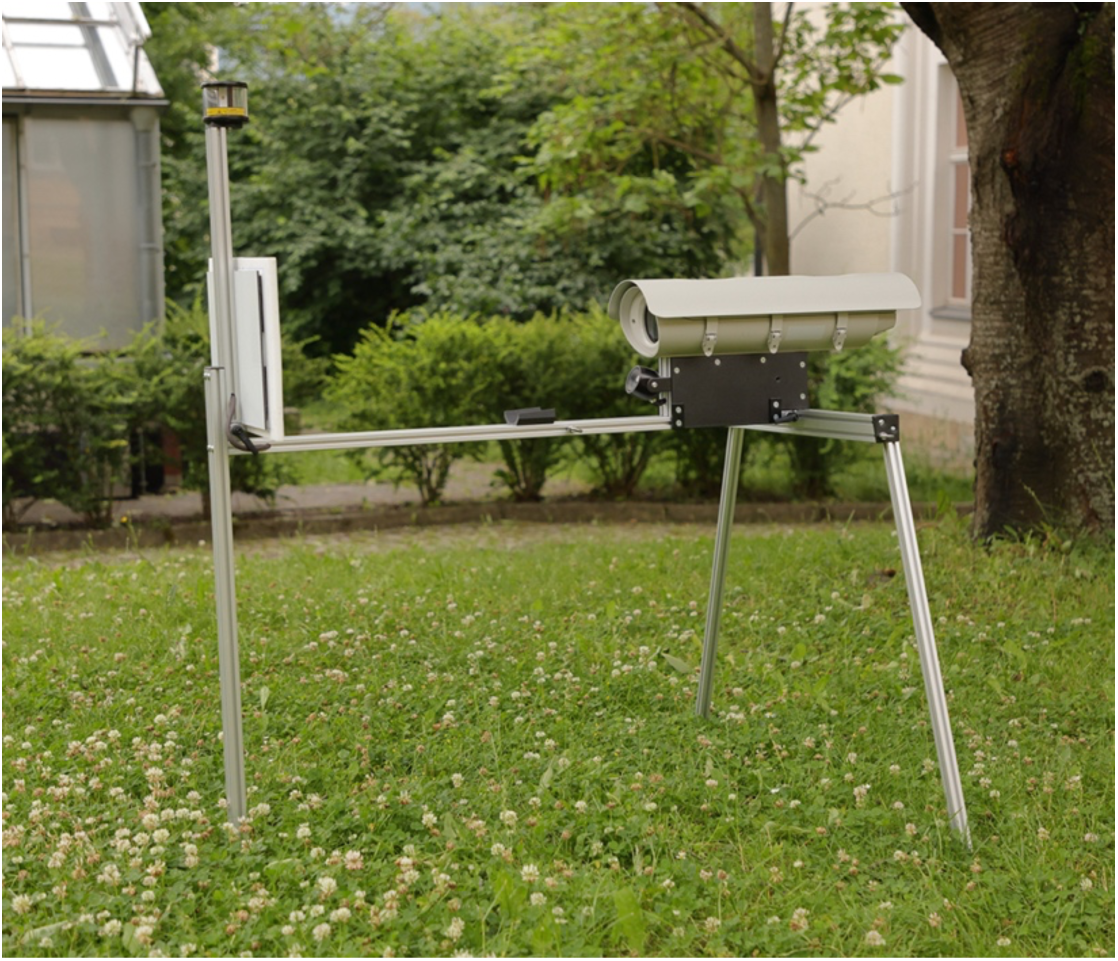
Prototype of the new CLT design in the LEPMON project (August 2024). Instead of a DSL camera, an industrial camera along with electronic parts are accommodated in a waterproof housing.

### Perspectives on AI for automated image data analysis

Modern approaches of AI and deep learning are nowadays able to solve complicated tasks like the identification of objects in real-world images and hence, they are also suited for analyzing image data recorded with CLTs automatically to reduce human labor. Species identification in images is a particular application of fine-grained image analysis and recognition (Wei et al., 2022), which deals with the automatic distinction of visually very similar looking categories at a subordinate level – in our case different species of nocturnal insects. A deep learning pipeline for automated visual moth monitoring has been presented by Korsch et al. (2021) for a CLT that was supposed to be part of a multisensor station for biodiversity monitoring within the AMMOD project (Wägele et al., 2022). Species classification by AI models can further be enhanced by exploiting images from the Internet as additional training examples, e.g., as shown by Böhlke et al. (2021). These AI approaches will be developed further within the LEPMON project to enable the automatic processing of large amounts of image data recorded by multiple CLTs. Besides that, other international project initiatives work on AI for nocturnal insect monitoring, e.g., Jain et al. (2024) and Padubidri et al. (2024).

### Conclusion and prospects

Our study provides important foundations for understanding benefits and limitations of automatic CLTs for nocturnal insects. We show that the method works well in principle, even though we still had some technical problems with the existing setup. In combination with AI, the method will provide a density and precision of data on nocturnal insects that cannot be achieved with any conventional method in future. In detail, CLTs seem to provide slightly different data than conventional traps. Further and regionally broader studies should follow so that the data from CLTs can be correctly interpreted. With further technological advances, CLTs could greatly help to better understand the global loss of insect species and develop measures against it.

## AUTHOR CONTRIBUTIONS

The study was designed by VH and GB. Fieldwork was carried out by VH. Data were statistically analyzed by VH and DB. The figures were prepared by all coauthors. The manuscript was written by VH and GB, with support from DB. PB contributed the paragraph on perspectives of AI.

## ACKNOWLEDGEMENTS

We express our thankfulness to Staff Foundation (Staff Stiftung Lemgo), which financed this work in a very uncomplicated and spontaneous manner with 4.500 € for the purchase of cameras and equipment. Christian Finke (Biological Station Paderborn/ Senne e.V.) helped us to select trap locations and with the funding application. We are also very grateful for the openness of Gerhard Brechmann and his family, who allowed us to collect the data on their land. Daniel Veit supported us a lot with the construction of the camera traps and Ralf Holzhauer helped with the construction of the weather shelters on site. Carsten Blum contributed the drone images of the study area and Egbert Friedrich checked the species list for plausibility.

## DATA AVAILABILITY STATEMENT

The data of this study are available in the supplementary material of this article.

## SUPPORTING INFORMATION

Additional supporting information can be found online in the Supporting Information section at the end of this article.

