## Supplementary Material for "Do camera light traps for moths provide similar data as conventional funnel light traps?"

**Tab. S1.** GPS data and elevation of the four trapping sites.

| | **GPS (decimal degree)** | **elevation (m a.s.l.)** |
|---|---|---|
| open habitat camera light trap (O CLT) | 51.89564, 8.66378 | 134 |
| open habitat funnel light trap (O FLT) | 51.89574, 8.66311 | 132 |
| forest habitat camera light trap (F CLT) | 51.89341, 8.66210 | 138 |
| forest habitat funnel light trap (F FLT) | 51.89312, 8.66113 | 140 |

**Tab. S2.** Light trapping night dates (funnel light traps and camera light traps operating in parallel).

| | |
|---|---|
| **April** | 1$^{st}$: 19.04.2023<br>2$^{nd}$: 24.04.2023 |
| **May** | 1$^{st}$: 15.05.2023<br>2$^{nd}$: 21.05.2023 |
| **June** | 1$^{st}$: 13.06.2023<br>2$^{nd}$: 18.06.2023 |
| **July** | 1$^{st}$: 14.07.2023<br>2$^{nd}$: 20.07.2023 |
| **August** | 1$^{st}$: 10.08.2023<br>2$^{nd}$: 20.08.2023 |
| **September** | 1$^{st}$: 16.09.2023<br>2$^{nd}$: 18.09.2023 |

**Tab. S3.** Individuals and species collected in the two habitats in this study.

| | Individuals | | | Species | | |
|---|---|---|---|---|---|---|
| **Family** | **forest** | **open habitat** | **total** | **forest** | **open habitat** | **total** |
| Drepanidae | 159 | 74 | 233 | 8 | 4 | 8 |
| Erebidae | 629 | 524 | 1153 | 25 | 27 | 30 |
| Geometridae | 1628 | 1120 | 2748 | 94 | 77 | 105 |
| Lasiocampidae | 8 | 8 | 16 | 3 | 3 | 4 |
| Limacodidae | 14 | 48 | 62 | 1 | 1 | 1 |
| Noctuidae | 374 | 503 | 877 | 48 | 60 | 72 |
| Nolidae | 10 | 15 | 25 | 3 | 5 | 5 |
| Notodontidae | 39 | 42 | 81 | 9 | 10 | 12 |
| Sphingidae | 7 | - | 7 | 1 | - | 1 |
| **Total** | **2868** | **2334** | **5202** | **192** | **187** | **238** |

**Table S4.** Richness and diversity in the 12:12 night comparison. Bold face: higher value.

| | Sp. obs | Chao ± SD | Jackknife ± SD | Bootstrap ± SD | Fisher's alpha | Shannon |
|---|---|---|---|---|---|---|
| CLT n=19 | 92 | **169.00** ± 30.21 | 140.32 ± 15.40 | 112.83 ± 8.53 | 34.62 | 3.92 |
| FLT n= 24 | **101** | 150.85 ± 18.15 | **151.79** ± 16.17 | **124.05** ± 9.31 | **51.70** | **4.19** |

**Tab. S5.** Species complexes in the study.

| Complex number | Included species |
|---|---|
| 1 | *Eilema caniola / complana* |
| 2 | *Eilema depressa / griseola / lurideola* |
| 3 | *Aplocera efformata /plagiata* |
| 4 | *Eupithecia abbreviata / exiguata / indigata / nanata / plumbeolata / pygmaeata / tantillaria / vulgata* |
| 5 | *Eupithecia icterata / pulchellata / linariata / succenturiata* |
| 6 | *Pasiphila chloerata / debiliata / rectangulata* |
| 7 | *Thera variata / britannica* |
| 8 | *Amphipyra pyramidea / berbera* |
| 9 | *Euxoa tritici / nigrofusca* |
| 10 | *Mesapamea secalis / secalella* |
| 11 | *Noctua janthe / janthina* |
| 12 | *Xestia ditrapezium/ triangulum* |

**Tab. S6.** Comparison of residence duration data, broken down by family.

| | total visibility time [min] | observation sequences | mean obs_seq duration [min] | median obs_seq duration [min] |
|---|---|---|---|---|
| Drepanidae | 4892 | 225 | 21.7 | 4 |
| Erebidae | 97264 | 3860 | 25.2 | 2 |
| Geometridae | 39652 | 2789 | 14.2 | 2 |
| Lasiocampidae | 32 | 14 | 2.3 | 2 |
| Limacodidae | 1754 | 57 | 30.8 | 6 |
| Noctuidae | 19818 | 841 | 23.6 | 2 |
| Nolidae | 896 | 20 | 44.8 | 2 |
| Notodontidae | 7532 | 61 | 123.5 | 18 |

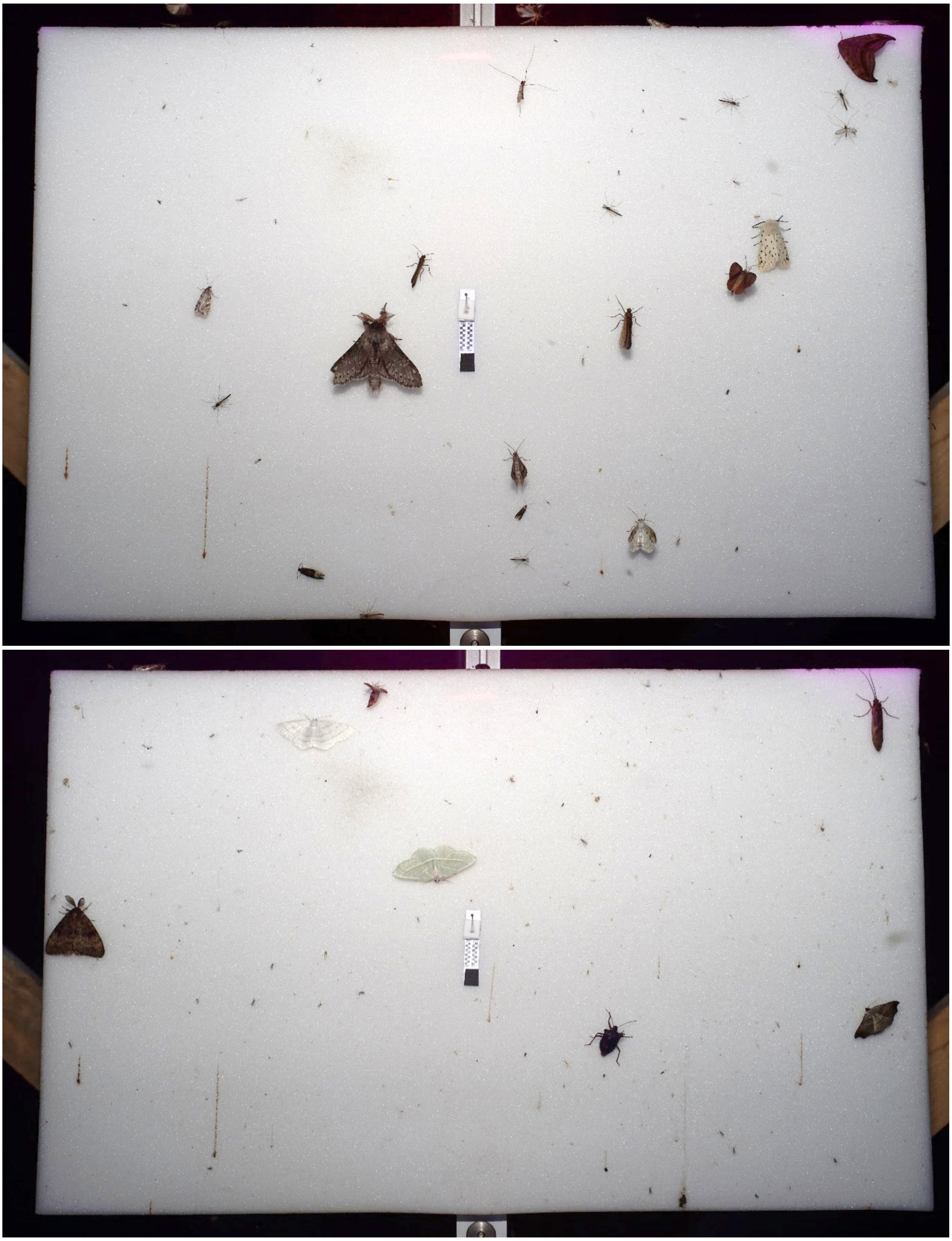

**Fig. S1.** Example of two of the resulting photos. A scale is attached in the center of the screen. Top: Macros that can be seen include *S. fagi, M. alternata/ notata, M. brunneata, L. temerata, S. lubricipeda, D. curvatula* and *L. marginata*, as well as 2 individuals with folded wings (probably *Thera* sp.). Furthermore, three micros are present, as well as some dipterans (mainly chironomids). 12/06/2023 01:31 a.m. – forest site. Bottom: *L. dispar, C. pusaria, C. margaritaria* and *L. flexula* as well as one Heteroptera and Trichoptera; 25/08/2023 00:14 a.m. – forest site.

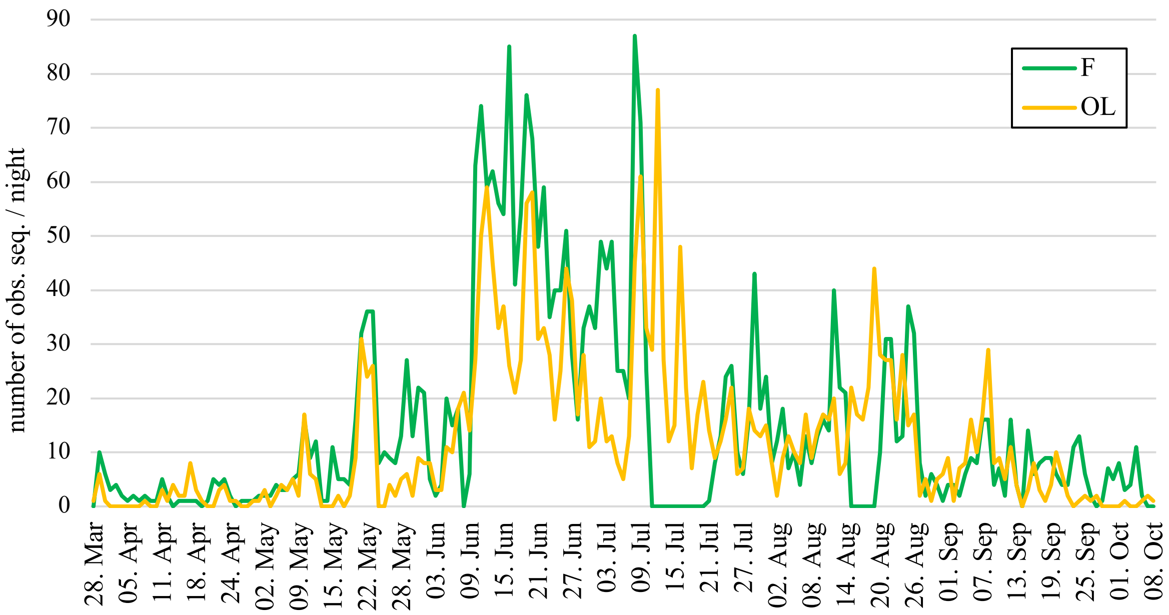


**Fig. S2.** Number of observation sequences per night broken down by the two habitat types in the course of the season. In sections where no observation sequences were made for several nights in a row, these were the nights where technical problems occurred, such as defective triggering of the flash of the camera. F = Forest, OL = Open habitat.

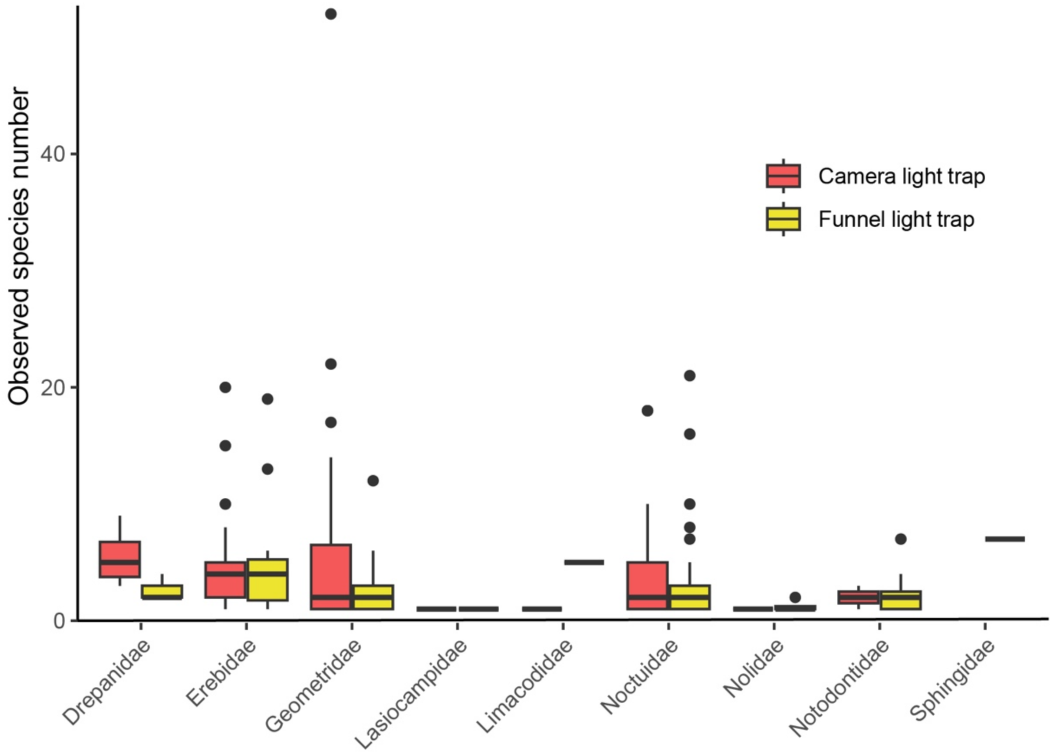


**Fig. S3.** Comparison of species richness observed in CLTs and FLTs of observed macromoth families.

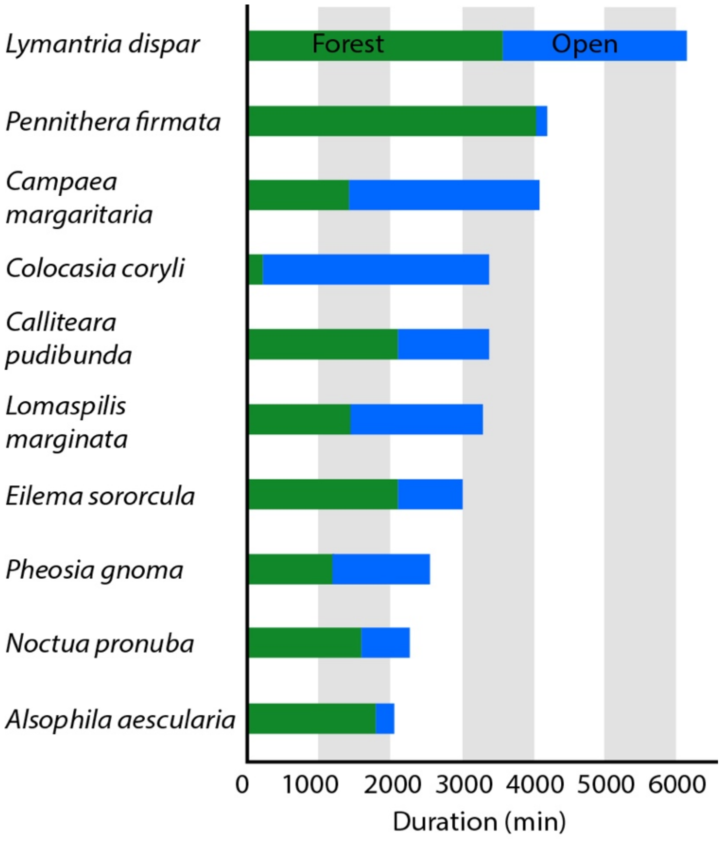


**Fig. S4.** Total (cumulative) visibility duration of ten moth species with longest observation time in CLTs (all 196 observation nights).
